# XenoSignal: Investigating Intra- and Inter-Species Ligand-Receptor Interactions Using AlphaFold3

**DOI:** 10.1101/2025.08.13.670200

**Authors:** Aly O. Abdelkareem, Courtney F. Hall, Athish Marutharaj, Kiran Narta, Theodore B. Verhey, A. Sorana Morrissy

## Abstract

Xenograft models, in which human tumors are implanted into mouse hosts, offer a unique platform to study tumor-microenvironment (TME) interactions. Ligand-receptor (LR) interactions are central to intercellular communication in this context, but most inference tools rely on single-species databases and assume conserved function across orthologues. To test whether this assumption holds given sequence and structural divergence, we developed XenoSignal, a computational framework that integrates AlphaFold3 structural predictions to assess the plausibility of intraspecies and interspecies LR interactions. The pipeline predicts whether human-mouse orthologues retain or lose their functional interaction, enabling construction of a high-confidence cross-species LR resource. XenoSignal facilitates the interpretation of tumor-TME crosstalk in xenografts and prioritizes LR pairs for experimental validation. This resource enhances our ability to dissect interspecies signaling, uncover mechanisms of immune evasion, and identify candidate therapeutic targets in translational cancer models.

## Introduction

Xenografts enable investigation of tumor progression, treatment responses, and complex intercellular interactions in a biologically relevant yet controlled context^1^. These interactions are largely mediated via ligand-receptor (LR) binding, whereby ligands from a sender cell bind to their cognate receptors on a receiver cell, initiating signalling cascades that regulate behaviour. Aberrant LR signalling is implicated in nearly every hallmark of disease^2^, and serves as an important focus of precision therapies.

Computational methods that infer LR interactions from bulk, single-cell, and spatial RNA sequencing data leverage curated databases of known LR pairs (e.g. STCase^3^, CellChat^4^, CellPhoneDB^5^, NICHES^6^). These are based on same-species LR interactions, and their direct use in xenograft settings^7^ assumes that interactions are functionally equivalent when either ligand or receptor is a different species. While this may be true for many LR pairs (e.g. human VEGF-A stimulates angiogenesis via mouse VEGF receptors^8^), substantial challenges to accuracy and interpretability arise when signaling molecules are not sufficiently conserved. In some cases, homologs may be entirely absent from one species (e.g. IL-8 in mice^9^), while in others, sequence or structural divergence can impact or abolish binding.

Although experimentally testing cross-species LR interactions is possible, the cost, low throughput, and reliance on specialized techniques or reagents^10^ limits this effort in practice. Instead, recent breakthroughs in AI-driven protein structure prediction could provide a scalable *in silico* alternative towards this goal. AlphaFold3 (AF3), for instance, can model protein-protein complexes at atomic resolution, accurately predicting binding interfaces^11^, and performs well even when interacting proteins originate from vastly different organisms^12^. We propose that a computational strategy to prioritize cross-species LR interactions for targeted experimental follow-up could be highly valuable by reducing the large search space to key interactions of high relevance.

Here, we systematically dock ligands and receptors within and between species, employing structure-based modelling with AF3. This workflow can in principle (i) account for subtle steric clashes at the binding interface, (ii) incorporate effects of multi-subunit assemblies and essential co-factors, and (iii) rank interactions by confidence score to prioritize those most or least likely to occur. The results and confidence metrics derived from the AF3 models are compatible with existing cell-cell communication analysis tools and ensure that the resulting knowledgebase - XenoSignal - is of immediate use to the research community. Ultimately, this resource bridges human and mouse signalling networks, advancing our understanding of tumor-TME crosstalk, and filling a knowledge gap in preclinical cancer research.

## Results

### A structure-based framework for systematic characterization of intra- and interspecies ligand-receptor interactions

We developed a structure-based computational framework to systematically evaluate human and mouse LR interactions within and across species using AF3^11^ **(Fig. 1a)**. A comprehensive database of 6,613 LR interactions among 2,821 unique genes from CellChatDB v1 and v2^4^ was assembled and extended to include mouse and human one-to-one orthologs (**Supplementary Table 1**). Addition of orthologous mouse-human and human-mouse versions of each LR pair generated a total of 13,546 unique inter- and intra-species interactions. To ensure compatibility with AF3, we applied preprocessing filters to exclude oversized complexes (max_combined_residues<5,000) and remove genes without both mouse and human protein sequences, yielding 13,283 unique candidate LR pairs. We evaluated 5,842 LR pairs (1183 genes) in which both partners are single-protein entities, to avoid complexity introduced by multi-protein assemblies **(Fig. 1b; Supplementary Table 2-3)**. These represented 240 unique pathways, of which 228 (95%) were shared between species (**Fig. 1c**).

**Figure 1.**
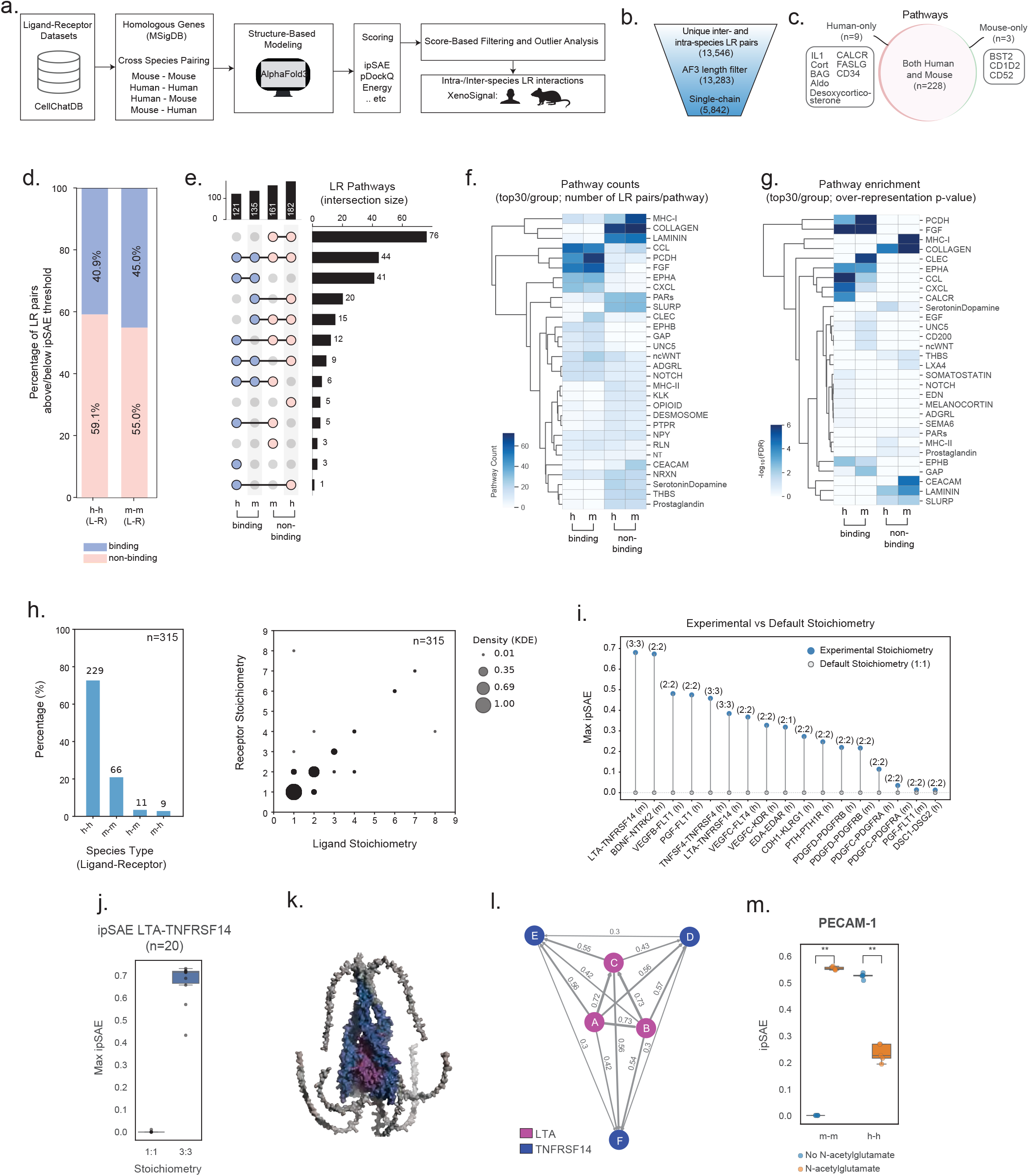
Schematic overview of the XenoSignal framework, dataset statistics, and intra-species ligand-receptor interaction analysis. --. **a**. Schematic overview of the XenoSignal computational pipeline. Starting from curated LR interaction datasets (CellChatDB), homologous ligand and receptor gene pairs were mapped across species using MSigDB ^29^, generating cross-species pairings (Mouse-Mouse, Mouse-Human, Human-Mouse, Human-Human). Each pair was structurally modeled using AlphaFold3 and scored using multiple metrics (ipSAE, pDockQ, energy, etc.). High-confidence predictions were filtered based on score thresholds, producing both intra- and inter-species LR interaction networks. **b**. Sequential filtering steps showing the number of unique inter- and intra-species LR pairs analyzed (13,546 total), the subset passing AlphaFold3 length constraints (13,283), and the final set of 5,842 high-confidence single-chain LR pairs used for downstream analyses. **c**. Venn diagram summarizing pathway-level overlaps for species-specific LR interactions among filtered single-chain pairs. A total of 228 pathways were conserved in both mouse and human, with 9 pathways unique to human and 3 unique to mouse. **d**. Distribution of ipSAE scores across intra-species LR pairs (Human-Human and Mouse-Mouse). Bars represent the proportion of binding (ipSAE > 0.17) and non-binding (ipSAE ≤ 0.17) predictions in each species. **e**. UpSet plot illustrating the overlap of pathway memberships across the four LR groups (binding and non-binding for human and mouse). The intersection sizes are shown as barplots. **f**. Heatmap quantifies the number of LR pairs per pathway across the four LR groups. The top 30 most frequent pathways (ranked by pathway count) are included (rows), and clustered based on LR composition profiles. Each cell indicates the count of unique ligand-receptor pairs per pathway. **g**. Heatmap of pathway enrichment significance (-log_10_FDR) across the same top 30 pathways, computed for each LR group. Color intensity reflects statistical significance of overrepresentation (FDR-capped at 6). **h**. Experimental LR stoichiometry data derived from PDB and Complex Portal. **Left:** Distribution of species type combinations (human-human, mouse-mouse, human-mouse, and mouse-human) among ligand-receptor pairs with available experimental structures (n = 315). Bar heights indicate percentages, with counts shown above each bar. **Right:** Receptor versus ligand stoichiometry shown as a dot plot. Each dot reflects the number of LR pairs with the given stoichiometry combination; KDE: kernel density estimate. **i**. Effect of stoichiometry on AlphaFold3-based binding predictions for selected intra-species LR complexes with ipSAE values of 0 under default stoichiometry (1:1). For each LR pair, the maximum ipSAE score is shown for both default (1:1) and experimentally-derived stoichiometries (e.g., 2:2, 3:3), with higher scores indicating stronger binding. Stoichiometry ratios are labeled on the plot, and human (h) or mouse (m) species is indicated in the label. **j**. Reproducibility of stoichiometry-specific predictions for the mouse Lta-Tnfrsf14 complex is shown as the distribution of maximum ipSAE values from 10 independent AlphaFold3 runs with different random seeds. **k**. 3D structural model of the mouse Lta-Tnfrsf14 complex in its native 3:3 stoichiometry. Chains are colored by identity (Lta in pink, Tnfrsf14 in dark blue), with clear visualization of the multimeric interface that drives complex formation. **l**. Inter-chain interaction network for the Lta-Tnfrsf14 3:3 model. Each node represents an individual chain, with ligand and receptor chains color-coded as in panel j. Edge thickness corresponds to AlphaFold3 interface confidence scores, and numeric values indicate pairwise interaction strengths, highlighting interface symmetry and multivalent engagement across subunits. **m**. Boxplots of ipSAE scores from multiple models are shown for human and mouse PECAM-1 models generated with (orange) and without (blue) PTMs. Significant changes are assessed with Mann-Whitney U test, where ** <= 0.01.

Quaternary structure of each LR pair was modelled with AF3. For each complex we computed a suite of structural confidence and interaction likelihood metrics, incorporating scoring strategies from AF3^11^, ipSAE^13^, PRODIGY^14^, and BindCraft^15^, including features such as ipSAE, iPTM, pDockQ^16,17^, predicted binding affinity, and other biophysical descriptors (see **Methods**). Together, these features constitute the XenoSignal knowledgebase (**Supplementary Table 4**). Clustering metrics identified seven groups **(Supplementary Fig. 1)**, including one enriched in features predictive of high-confidence binders (cluster4). We selected ipSAE as our primary metric based on its bimodal distribution suggestive of strong discriminative power in our data (**Supplementary Fig. 2**), and in agreement with previous benchmarks^13^. We classified the 5,842 inter and intra-species LR pairs into candidate binders and non-binders (ipSAE threshold=0.17; **Fig. 1d)**, finding that 41% of human (n=560), and 45% of mouse (n=660) same-species LR pairs were predicted to bind (**Fig. 1e**). Notably, binding versus non-binding status was highly consistent among species for 76.2% of pathways (**Fig. 1e**), and among classes of pathways (e.g. poor modelling of collagens and laminins in both species; **Fig. 1f-g**). These observations strongly indicate that additional context-dependent variables (e.g. cofactors, stoichiometry, post-translational modifications (PTMs)) are necessary to improve predictions for non-binders. Moreover, given the general conservation of non-binders among species, our results suggest that variables influencing LR interaction predictions are likely to be conserved cross-species. If substantiated, then identifying the contribution of a given variable to binding in one species could improve structural modelling for the other.

We speculated that stoichiometry contributes significantly in this way, as LR subunit counts are not experimentally available for most complexes. Of 315 LR complexes in our dataset with available experimental structures, 43.2% (136) had experimental stoichiometry ratios >1:1 (**Supplementary Table 5**; **Fig 1h**). Extrapolating this to the full list of 5,842 LR pairs, we anticipate that 2523 are incorrectly modelled with the default 1:1 ratio used in AF3, affecting the accuracy of many predictions. To explore the impact of non-default stoichiometry on AF3 predictions, we selected a subset of 17 non-binding LR pairs (ipSAE=0) with experimental data, and found that incorporating experimental stoichiometries boosted models to the binding range in 76.5% of cases (13/17 pairs; ipSAE>0.17; **Fig 1i-**), and often in both species (e.g. LTA-TNFRSF14;-**Fig. 1i-l**). In addition to stoichiometry, we anticipated that predictions may benefit from inclusion of PTMs, and we identified multiple non-binders in this category (**Supplementary Table 6**). For instance, PECAM-1 (CD31), binding in mice is dependent on glycan modifications^18^. Incorporating these PTMs into the AF3 model markedly increased ipSAE scores for mouse but not human PECAM-1 **(Fig. 1m**), in line with experimental findings(**Supplementary Table 7**).

### Disrupted Interspecies Binding

We quantified significant disruptions in cross-species binding using the change (delta) in ipSAE scores when either the receptor or ligand was set to the other species (**Fig. 2a-b**). Although a strong relationship emerged between sequence conservation and binding (r=0.39, pval=1.6e-12), many highly conserved LR pairs had disrupted interspecies binding (e.g. 53 LR pairs with homology>80%). We manually curated a set of outliers using literature evidence (n=46; see **Methods**; **Figure 2a-c**), finding full or partial support for lowered interspecies binding in several cases, including TNFSF9 (4-1BBL)-TNFRSF9 (4-1BB)^19,20^ and TNFSF18 (GITRL)-TNFRSF18 (GITR)^21^ (**Fig. 2d**; **Supplementary Table 6-8**). In these specific complexes stoichiometry ratios differences between species are known to impact binding. Re-modelling both pairs with species-specific stoichiometry maintained the identified disruptions in binding (**Supplementary Table 6-8; Fig. 2e**).

**Figure 2.**
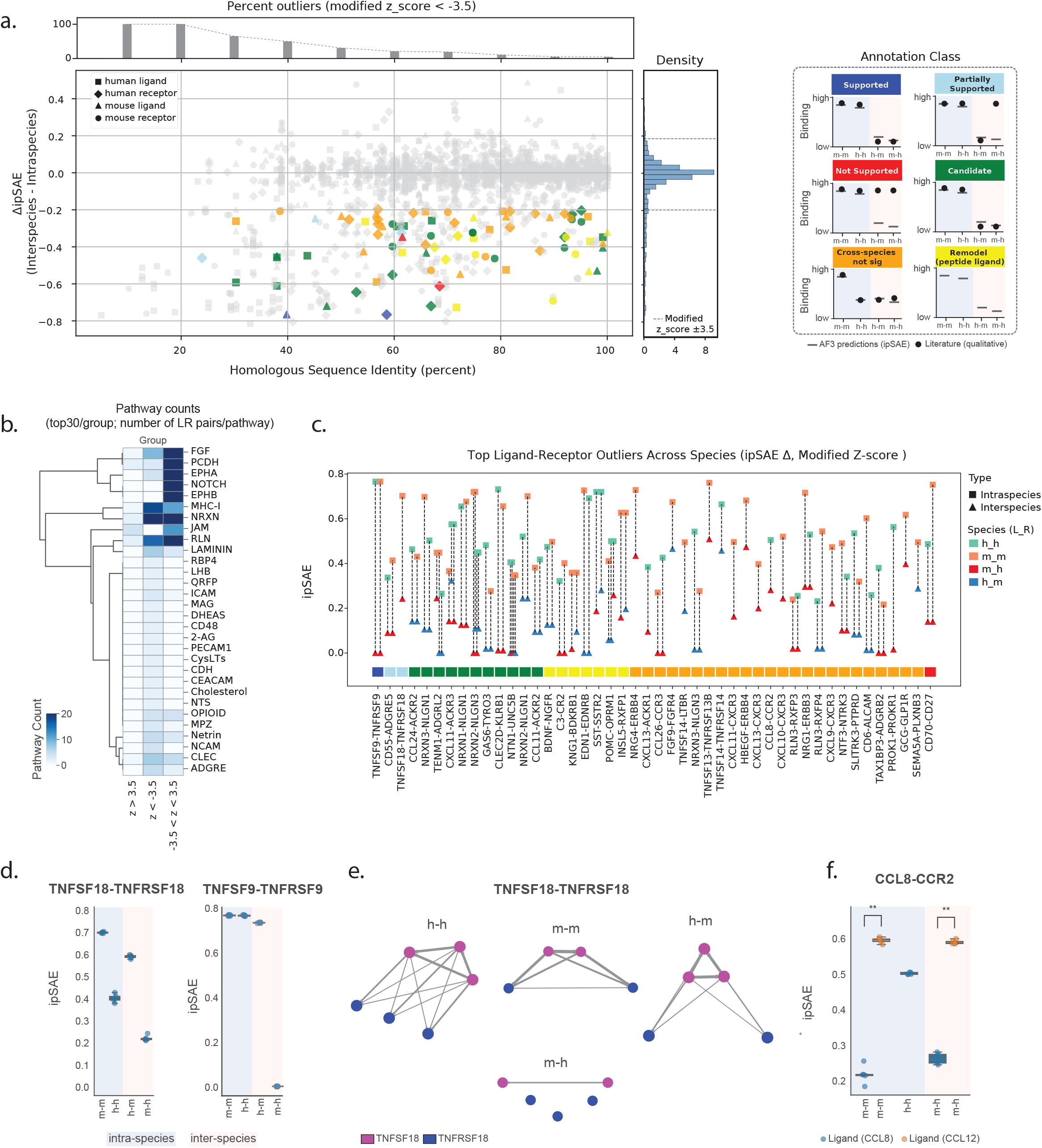
Inter-species analysis of ligand-receptor interactions using XenoSignal. **a**. The relationship between amino acid sequence identity and change in predicted binding scores across species is shown as a scatterplot. Each point represents an LR pair. The change in ipSAE score (ΔipSAE) is calculated by subtracting each intraspecies model ipSAE value from the interspecies value. (y-axis). Negative ΔipSAE values indicate reduced predicted binding in the cross-species context. Percent identity (x-axis) represents the minimum amino acid sequence identity between human and mouse orthologs. Colored points highlight significant outliers (modified z-score < -3.5) with predicted disruption in interspecies binding. The histogram panel above the scatter plot quantifies the proportion of negative outliers across quantiles of sequence identity. The density plot in the right panel illustrates the overall distribution of ΔipSAE scores, with a dashed line marking the outlier threshold. A classification legend defines the curated annotation categories corresponding to the colored points. “Supported” (blue) indicates LR pairs with concordant ipSAE score and literature evidence; “Partially Supported” (green) includes cases where the literature supports binding disruptions in one interspecies context (either m-h or h-m); “Not Supported” (red) reflects a contradiction with literature; “Candidates” (yellow) do not have literature support; “Cross-species not significant” (orange) are LR pairs where drops in interspecies binding scores are not much lower than in either m-m or h-h; “Remodel (peptide ligand)” (yellow) represent LR pairs where the ligand undergoes proteolytic cleavage that is not captured in current models; and (Grey) are additional disrupted LR pairs without curation. **b**. Heatmap showing the number of LR pairs per pathway across three distribution groups (z > 3.5, z < -3.5, and -3.5 ≤ z ≤ 3.5). The top 30 pathways with the most significant FDR values (ranked by pathway count) are displayed as rows and clustered based on LR composition profiles. Each cell represents the number of unique LR pairs for a given pathway, capped at a maximum of 20 counts. **c**. Top cross-species outliers ranked by annotation class and ΔipSAE. Each vertical line connects points representing ipSAE scores for an LR pair modeled in intraspecies (square) and interspecies (triangle) contexts. Interactions with intraspecies binding (ipSAE>0.17) and disrupted interspecies binding (ΔipSAE ≤ -0.2; modified z-score < -3.5) are included. Points are colored by LR species configuration (human-human, human-mouse, mouse-human, mouse-mouse), and the colored bar below reflects the same annotation classes as in panel (a). **d**. Structural and functional resolution of selected outlier cases. ipSAE values for five predicted AlphaFold3 (AF3) models per run, shown as boxplots, for either default (1:1) or experimentally derived stoichiometries. **e**. Structural representations of the TNFSF18-TNFRSF18 complex modelled under different conditions. Nodes represent ligand (magenta) and receptor (marine) chains, with edges indicating predicted physical interactions. Edge thickness corresponds to AlphaFold3 interface confidence scores. Absent edges indicate no interaction or ipSAE< 0.17. **f**. ipSAE values for five AF3 models per run for two alternative ligand orthologs (CCL8 and CCL12) interacting with CCR2, shown as boxplots.

Several of the 46 curated LR pairs were peptide ligands that undergo proteolytic processing into active forms (e.g. SST-SSTR2 and POMC-OPRM1^22^). Since ligands are currently modelled as whole proteins, this LR class may thus require re-modeling to incorporate proteolysis-specific features. There were several additional LR pairs without experimental data, not known to be processed into peptide ligands, and lacking annotated PTMs. These constitute a shortlist of candidates with disrupted interspecies binding that warrant further experimental investigation.

Finally, we highlight CCL8-CCR2, a case where AF3 predictions show poor binding of the annotated mouse CCL8 ortholog to mouse or human CCR2 receptors (**Fig. 2f**). This prediction informs ongoing debate as to whether *Ccl12*, and not *Ccl8*, is the functional mouse ortholog for human *Ccl8*^23^. By modelling both ligands, we demonstrate strong AF3 support for functional interactions between mouse CCL12 and both mouse and human CCR2 (**Fig. 2f**). This finding suggests that our AF3 modelling strategy can help resolve ambiguity for genes with complex orthologous relationships^24^.

## Discussion

Our structure-based computational framework, XenoSignal, offers a scalable approach for assessing interspecies cellular crosstalk based on modelling of ligand-receptor complexes. We demonstrate that most LR complexes retain binding between mouse and human orthologs, indicating highly conserved functional interactions in accord with the strong assumptions underpinning xenograft data interpretation. However, we also identify an important subset of genes that deviate from this trend, including known and novel cases of species-specific divergence. These require bespoke assessments and experimental follow-up to ensure appropriate interpretation of functional relevance in preclinical studies.

Several limitations of our current implementation highlight avenues for future development. First, stoichiometry emerged as a major contributor to prediction accuracy, and we anticipate that emerging methods to predict subunit counts will significantly boost modelling success for many complexes^25^. In addition, incorporation of heteromeric complexes, complexes larger than 5000 residues, addition of cofactors, PTMs, and consideration of isoforms^26,27^, could all extend our findings. Finally, the field lacks standardized benchmarks defining how AF3 scores correspond to biological binding in different contexts. We anticipate this challenge will be addressed with the rapid growth of structural modelling-based binder discovery efforts that incorporate experimental validations^15^.

In summary, XenoSignal establishes a compelling proof-of concept framework for scalable cross-species LR binding prediction that accelerates rapid hypothesis generation and screening of cellular interactions. XenoSignal is compatible with existing LR inference workflows, and readily extendable to incorporate other model species.

## Methods

### Liand Receptor Dataset Generation

We developed a comprehensive cross-species LR interaction dataset. The generation pipeline consisted of five major stages: database integration, ortholog mapping, LR interaction expansion, protein sequence retrieval, and preparation for AlphaFold 3^11^ submission.

#### 1. Database Integration

We began with the CellChatDB v2^4^, which includes curated intra-species ligand-receptor interactions with support for complexes and cofactors. Specifically, the human dataset comprises 3,234 interactions, including 338 protein complexes and 32 cofactors, while the mouse dataset includes 3,379 interactions with 337 complexes and 33 cofactors. Merging the human-human and mouse-mouse interactions resulted in 6,613 total LR interactions, integrating 3,914 interactions from CellChatDB v1 and 2,699 newly added interactions from v2.

#### 2. Ortholog Mapping

To enable human-mouse and mouse-human cross-species mapping, we constructed an ortholog dictionary by combining three resources:

- NICHES ^28^ ortholog dataset: includes 18,758 mouse-to-human and 18,308 human-to-mouse gene mappings.
- MSigDB v7.5.1^29^ : contributes 101,764 mouse-to-human and 18,323 human-to-mouse orthologs. After removing duplicates, we identified 83,996 unique mouse-to-human mappings that were not captured by NICHES.
- Manual curation: we identified and added 87 ortholog pairs (human-mouse or mouse-human) that were missing from both datasets.

#### 3. Mapping Ligand-Receptor Interactions Across Species

We applied the ortholog dictionary to the 6,613 original intra-species interactions to generate interspecies LR pairs. For each interaction, if both ligand and receptor genes had corresponding orthologs in the other species, we generated two additional interactions: human ligand-mouse receptor and mouse ligand-human receptor. This expansion process yielded 19,839 total LR interactions. Of these, 6,294 were duplicates arising from bidirectional mapping (i.e., the same interaction generated from both human-mouse and mouse-human permutations). After removing these redundancies, we retained 13,283 unique LR interactions.

#### 4. Protein Sequence Retrieval

We identified 2,875 unique genes involved in the 13,283 LR interactions (**Supplementary Table 9)**. For each gene, we queried the NCBI Protein database to retrieve the amino acid sequence corresponding to its canonical isoform. Retrieved sequences were stored in a custom CellChat protein dataset. A total of 60 genes lacked protein sequences in NCBI, resulting in the exclusion of 181 LR interactions from downstream structural modelling.

#### 5. Preparation for AlphaFold 3 Submission

To prepare for structural modelling using AlphaFold 3, we considered several submission constraints: (i) each JSON input file can contain up to 100 jobs, (ii) a daily limit of 20 job submissions is enforced, and (iii) the combined amino acid length of ligand and receptor must not exceed 5,000 residues. We calculated the total sequence length for each LR pair and excluded interactions exceeding the length threshold. We also generated a histogram of sequence lengths to visualize the distribution across interactions and flagged prioritized interactions to be included in the earliest AlphaFold 3 submissions. After filtering for available protein sequences, removing duplicates, and applying the amino acid length constraint, the final dataset comprised 13,283 unique LR interactions suitable for AlphaFold 3-based structural prediction.

### Calculation of Interaction Features

Following AlphaFold 3 structural predictions, we computed a comprehensive set of interaction features to quantitatively assess the quality, stability, and biophysical characteristics of the predicted ligand-receptor complexes (**Supplementary Table 2-3**). These metrics derived using established methods, provide a multidimensional evaluation of docking accuracy and binding efficiency. Collectively, these detailed features enable a rigorous assessment of both structural quality and biophysical properties for each complex, thereby facilitating the prioritization of interactions for further experimental validation and mechanistic studies.

#### 1. Predicted Docking Quality and Accuracy

- **pDockQ** (pDockQ): A score estimating docking confidence based on interface geometry and surface complementarity. Higher values reflect more stable and likely biologically relevant interactions^16^.
- **pDockQ2** (pDockQ2): A docking score derived from pairwise predicted aligned error (PAE), offering improved discrimination between high- and low-confidence models^17^.
- **LIS** (LIS): The Local Interaction Score, computed from transformed PAE values, highlights residue-level interaction confidence^30^.

#### 2. IpSAE Features^13^

- **ipSAE** (ipSAE): interaction prediction Score from Aligned Error, computed using PAE and residues from chain 2 under a PAE cutoff.
- **ipSAE_chn** (ipSAE_d0chn): ipSAE using a distance reference d0d_0d0 equal to the sum of chain lengths.
- **ipSAE_dom** (ipSAE_d0dom): ipSAE using d0d_0d0 as the number of residues with interchain PAE below the threshold.
- **n0_res** (n0res): The number of residues used to define d0d_0d0 in ipSAE calculation.
- **n0_chn** (n0chn): The d0d_0d0 chain-length reference used in ipSAE_chn.
- **n0_dom** (n0dom): The d0d_0d0 domain-level reference for ipSAE_dom.
- **nres1** (nres1) and **nres2** (nres2): Number of residues in each chain under the PAE cutoff and with an interacting partner.
- **dist1** (dist1) and **dist2** (dist2): Number of residues in each chain under both distance and PAE thresholds.

#### 3. AlphaFold Structural Confidence^11,31^

- **ipTM_af** (ipTM_af): AlphaFold-predicted TM-score for the full complex, obtained from model output.
- **ipTM_chn** (ipTM_d0chn): TM-score computed from the PAE matrix, normalized using the sum of chain lengths.
- **frac_disordered** (af3_fraction_disordered): Proportion of residues predicted to be disordered by AlphaFold3.
- **af3_iptm** (af3_iptm): ipTM metric from AlphaFold3, reporting predicted inter-chain TM-score.
- **af3_ptm** (af3_ptm): Predicted TM-score from AlphaFold3, often used for assessing monomer fold quality.
- **pLDDT_mean** (i_pLDDT): Mean predicted Local Distance Difference Test score over interface residues.
- **pLDDT_ss** (ss_pLDDT): pLDDT values restricted to residues in secondary structure elements (e.g., helices, strands).

#### 4. Intermolecular Contacts^14^

- **n_contacts** (Number of Intermolecular Contacts): Total residue-residue contacts across the interface.
- **cc_contacts** (Charged-Charged Contacts): Number of contacts between charged residues.
- **cp_contacts** (Charged-Polar Contacts): Charged-polar contact count.
- **ca_contacts** (Charged-Apolar Contacts): Charged-apolar contact count.
- **pp_contacts** (Polar-Polar Contacts): Contacts between polar residues.
- **ap_contacts** (Apolar-Polar Contacts): Apolar-polar residue interactions.
- **aa_contacts** (Apolar-Apolar Contacts): Apolar-apolar contact count.

#### 5. Surface Composition and Energetics^14^

- **apolar_NIS_pct** (Percentage of Apolar NIS Residues): Fraction of apolar residues on the Non-Interacting Surface.
- **charged_NIS_pct** (Percentage of Charged NIS Residues): Fraction of charged residues on the Non-Interacting Surface.
- **pred_affinity** (Predicted Binding Affinity (kcal/mol)): Binding free energy estimated in kcal/mol.
- **pred_Kd** (Predicted Dissociation Constant (Kd, M)): Estimated dissociation constant at 25°C, indicating binding strength.

#### 6. Energetic and Structural Metrics^15^

- **unrelaxed_clashes** (Unrelaxed_Clashes): Number of atomic clashes in unrelaxed AlphaFold models.
- **binder_energy** (Binder_Energy_Score): Total interface energy, often used as a design or docking quality indicator.
- **surf_hydrophobicity** (Surface_Hydrophobicity): Average hydrophobicity of exposed surface residues.
- **shape_comp** (ShapeComplementarity): Geometric complementarity score of interacting surfaces.
- **packstat** (PackStat): Packing density at the interface, with higher values indicating better residue fitting.
- **dG** (dG): Predicted Gibbs free energy of binding.
- **dSASA** (dSASA): Solvent-accessible surface area buried upon binding.
- **dG_dSASA** (dG/dSASA): Binding energy normalized by surface area buried, measuring efficiency.

#### 7. Interface Structural Features

- **intf_SASA_pct** (Interface_SASA_%): Proportion of complex surface area buried upon interface formation.
- **intf_hydrophobicity** (Interface_Hydrophobicity): Hydrophobicity index for residues at the binding interface.
- **n_intf_res** (n_InterfaceResidues): Number of residues forming the interface.
- **n_intf_Hbonds** (n_InterfaceHbonds): Number of hydrogen bonds across the interface.
- **intf_Hbonds_pct** (InterfaceHbondsPercentage): Fraction of interface residues engaged in hydrogen bonding.
- **n_unsat_Hbonds** (n_InterfaceUnsatHbonds): Count of unsatisfied hydrogen bonds at the interface.
- **unsat_Hbonds_pct** (InterfaceUnsatHbondsPercentage): Proportion of H-bonds not forming stable pairs.
- **intf_helix_pct** (Interface_Helix%): Fraction of interface residues in α-helices.
- **intf_beta_pct** (Interface_BetaSheet%): Fraction in β-sheet conformation.
- **intf_loop_pct** (Interface_Loop%): Fraction in loops or coils.
- **binder_helix_pct** (Binder_Helix%): Overall helical content of the binder chain.
- **binder_beta_pct** (Binder_BetaSheet%): β-sheet content of the binder chain.
- **binder_loop_pct** (Binder_Loop%): Loop content in the binder.

### Clustering of Binding Features

To uncover patterns in protein interaction features, we computed a pairwise distance matrix from the **absolute Pearson correlation** of the selected features. The resulting square matrix was converted to condensed form with SciPy’s squareform function^32^, and hierarchical clustering was performed using the average linkage method. A fixed distance threshold of t=0.75 was applied to define discrete clusters with the criterion=‘distance’ parameter. This approach partitioned the data into seven distinct clusters, representing interaction classes with shared properties.

### Determination of a Binding Threshold based on ipSAE Distribution

To distinguish weak from strong ligand-receptor interactions, we identified a data-driven threshold based on the bimodal distribution of ipSAE values. Kernel density estimation (KDE) was applied to smooth the distribution, and local maxima and minima were identified using SciPy’s find_peaks^32^. When two peaks were detected, the lowest point (valley) between them was selected as the threshold; otherwise, the midpoint between peaks or the global minimum was used. This analysis yielded a threshold of 0.17, which we used to classify interactions into non-binders (ipSAE ≤ 0.17) and binders (ipSAE > 0.17).

### Protein Homology Analysis

To assess protein homology, we compared amino acid sequences of ligand or receptor proteins either within the same species or across species when common ligands or receptors were identified. Protein sequences were retrieved from the NCBI Entrez Protein database using accession IDs, and global pairwise sequence alignments were performed using Biopython’s pairwise2 module^33^ with identity scoring (globalxx). For each protein pair, the best alignment was selected, and the percentage identity was calculated as the number of identical residues divided by the length of the longer sequence, serving as a measure of sequence-level homology.

### 3D Structure Modeling and ipSAE Graph Construction

To visualize ligand-receptor complexes, we rendered 3D structures from Crystallographic Information Files (CIFs) using PyMOL (Version 1.2r3pre, Schrödinger, LLC.). Structural representations followed a standardized rendering protocol to enhance surface detail and interfacial visibility. Specifically, surfaces were displayed with 80% transparency, high-resolution rendering enabled, and two-sided lighting activated. Additional enhancements included shadowing, plastic lighting style, and modernized rendering settings to maximize clarity and depth perception.

Using structural models involving interactions among more than two protein chains, we first computed ipSAE scores between all chain pairs, then visualized these interactions as directed graphs. For each pair of chains, the maximum ipSAE value across all interface orientations was retained to represent the most substantial contact. A directed edge was drawn from one chain to another if the ipSAE exceeded a defined threshold (≥ 0.17), with edge thickness proportional to the interaction strength. Nodes in the graph represent individual chains and were colored according to their functional classification (e.g., ligand or receptor) based on user-defined annotations. Graphs were arranged using the Kamada-Kawai force-directed layout to improve visual clarity. This graph-based representation provided an interpretable summary of the inter-chain interaction topology, highlighting dominant interfaces within multimeric protein assemblies.

### Cross-Species Outlier Detection Based on ipSAE Shifts

To identify ligand-receptor pairs exhibiting unexpected shifts in interactions between cross-species and same-species comparisons, we focused on interactions involving either a common ligand or receptor shared between human (h) and mouse (m). For each LR pair, we computed the change in interfacial exposure between interspecies and intraspecies complexes as (**Supplementary Table 10**):

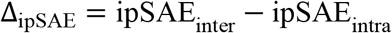

The distribution of Δ_ipSAE_ values was visualized using a kernel density plot. To robustly detect outliers, we applied a modified z-score filter using the median and median absolute deviation (MAD), defined as:

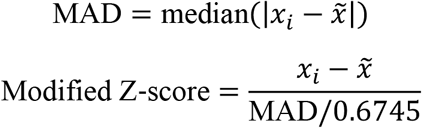

Outliers representing disrupted interspecies binding were defined as pairs with modified z-scores below −3.5. We note that outliers with modified z-scores above +3.5 could represent cases where (1) AF3 performs less well intra-species relative to inter-species, or (2) the cross-species ortholog annotated in the LR databases is incorrect.

To avoid spurious hits driven by poor interaction modelling in the same species, we excluded any pair where the intraspecies ipSAE score was <0.17.

To further assess the absence of meaningful cross-species binding, we calculated the average intra- and interspecies ipSAE values for each pair:

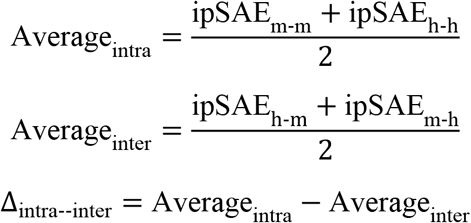

Pairs with Δ_intra--inter_ < 0.13 were classified as exhibiting no significant cross-species binding. Together, these criteria enabled the identification of both structural outliers and potential species-specific interaction failures.

### Pathway Enrichment Analysis

To compare functional signatures of ligand-receptor interactions within the same species, we performed hypergeometric pathway overrepresentation tests on four groups—Human Binders, Mouse Binders, Human Non-binders, and Mouse Non-binders—using the top LR as background. For each group, we extracted FDR-adjusted q-values and in-group gene counts per pathway and assembled two matrices: one of -log10(FDR) scores and one of gene counts. We then selected the 30 pathways with the highest peak significance across groups.

### Retrieval of Experimental Structures and Stoichiometry Ratios for LR Pairs

To obtain experimental stoichiometry ratios for LR interactions, UniProt accession codes for each ligand and receptor—including synonymous gene names—were retrieved using the UniProt ^34^ and BioNet ^35^ APIs, then filtered to retain only human and mouse entries, with additional manual curation. PDB entries for each ligand and receptor were then identified via the RCSB ^36^ search API to locate experimental structures, with selected synonym mappings validated using UniProt’s Retrieve/ID mapping tool. Stoichiometry ratios were initially extracted from RCSB’s PISA data; however, inconsistencies in chain ID assignments required remapping entities to assemblies, accounting for author-assigned chain IDs, and recalculating accurate ratios. For each PDB entry, ligand and receptor stoichiometries, accessions, organism, sequence coverage, length range, and post-translational modifications were recorded. Stoichiometry ratios (L:R) were consolidated across assemblies, yielding 806 PDB records. After filtering redundant entries and retaining those with the highest L:R ratio per interaction, 307 non-redundant experimental structures were obtained. An additional eight stoichiometry records were incorporated from Complex Portal for interactions lacking structural data, resulting in a final set of 315 LR stoichiometry records used in the analysis (**Supplementary Table 5**). This dataset comprises 72.7% human-human (h-h; *n* = 220), 21.0% mouse-mouse (m-m; *n* = 66), 3.5% human-mouse (h-m; *n* = 11), and 2.9% mouse-human (m-h; *n* = 9) interactions. Notably, 179 records (56.8%) exhibit a 1:1 LR stoichiometry ratio.

## Reporting summary

Complete.

## Data Availability

All raw AlphaFold3 job submission files, including input sequences and random seeds used to generate the structural predictions in this study, are available at: https://github.com/MorrissyLab/XenoSignal/Data. These files are provided to ensure full reproducibility of the predicted structures analyzed in this work.

## Code Availability

The XenoSignal framework is available as open-source software. All code used to perform analyses and generate figures in this study is publicly accessible at: https://github.com/MorrissyLab/XenoSignal.

## Supplementary Information

Supplementary Figures 1-2 and Supplementary Tables 1-10 included in the main submission file.

## Author Information

### Contributions

Conceptualization, AA, ASM; Methodology, AA, ASM, CFH; Investigation, AA, CFH, TBV, AM, KN, ASM; Data curation, AA, CFH; Writing-Original draft, AA, ASM; Writing-Review & Editing, AA, ASM; Funding acquisition, ASM; Supervision, ASM.

## Funding Acknowledgements

ASM was supported by a Canadian Institutes of Health Research (CIHR) Operating Grant [grant number 400678] and holds a Canada Research Chair (CRC) Tier 2 in Precision Oncology. AA was supported by the Alberta Children’s Hospital Research Institute Graduate Scholarship.

## Acknowledgements

We gratefully acknowledge the indispensable contributions of Mohammed Korayem, Ahmad Hafez, Hossam Emara, Tarek Talaat, Muhammad Hammam, Gamal Alzeer, and Yasser Hendwai, who played a crucial role in submitting large-scale AlphaFold3 prediction jobs during its pre-release phase, despite the strict limit of 20 jobs per Google account per day. By efficiently coordinating submissions across multiple accounts for weeks, they enabled the extensive computational throughput necessary for this study.

## Supplementary Figure Legends

**Supplementary Figure 1.**
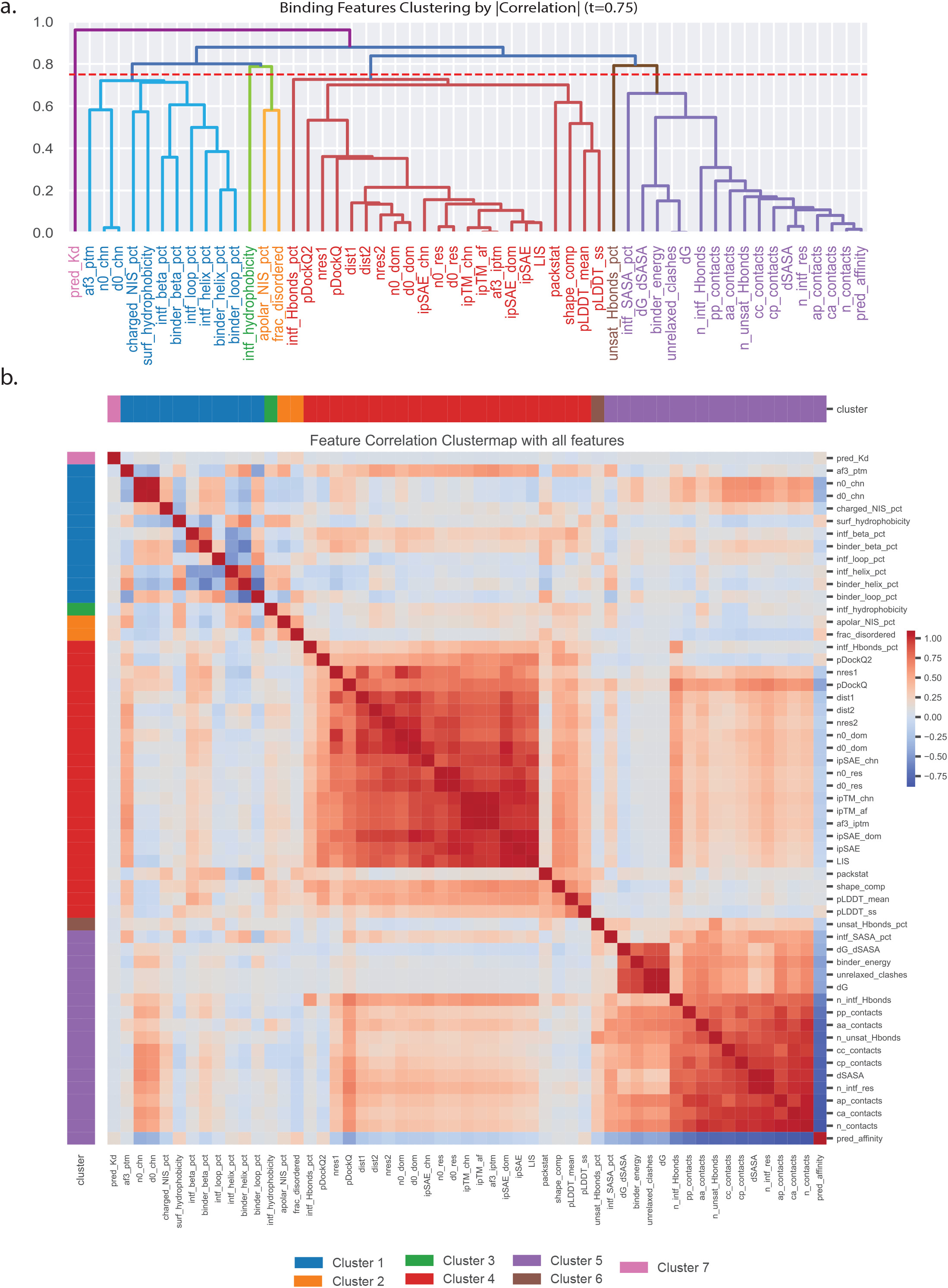
Hierarchical clustering of structural and binding features for ligand-receptor complex evaluation. **a**. Identification of key structural features for binding discrimination. Dendrogram showing the hierarchical clustering of binding and structural metrics (n = 23) organizes features into seven clusters, highlighting relationships between features and guiding the selection of ipSAE as the primary discrimination score. **b**. Pairwise correlation matrix of all computed features used to assess ligand-receptor interactions. Features are organized by hierarchical clustering (threshold = 0.75), delineating clusters of highly correlated metrics that capture related structural or biophysical properties. Color bars indicate cluster assignments. The clustermap reveals groups of redundant features as well as distinct, non-redundant metrics, guiding the selection of representative features for downstream analysis and benchmarking.

## Supplementary Figure 2

**Supplementary Figure 2.**
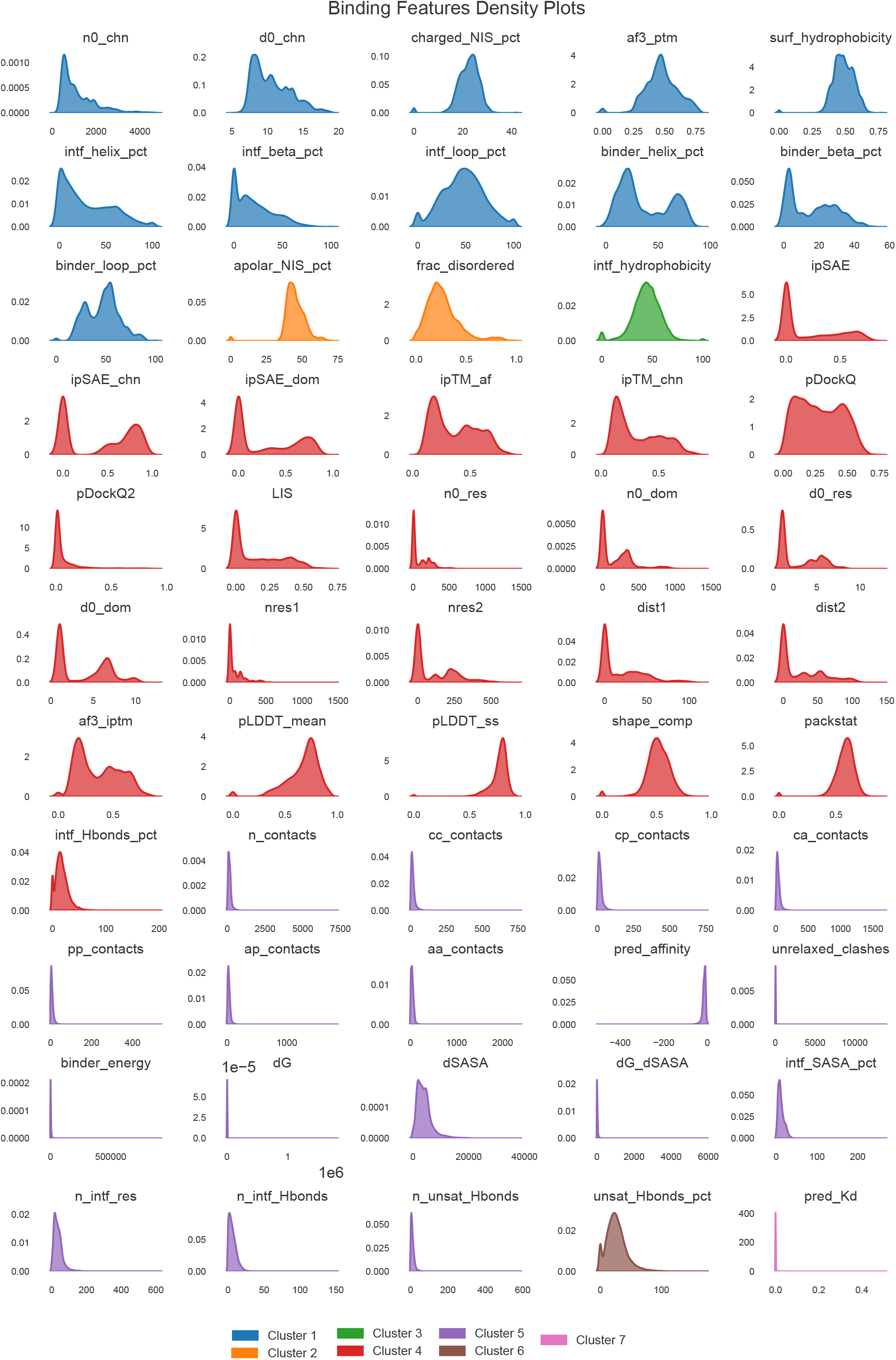
Density distributions of binding features grouped by cluster. Kernel density plots of all structural and binding features used for ligand-receptor complex evaluation, grouped and color-coded according to their cluster assignments (seven clusters, as defined by hierarchical clustering in Supplementary Fig. 1). Each plot displays the distribution of feature values across the dataset. Features within the same cluster exhibit similar distribution patterns—such as unimodal or bimodal shapes—highlighting shared biophysical or functional properties. This clustering aids in feature reduction and the identification of representative metrics for downstream analyses.

## Description of Additional Supplementary Files

Supplementary Tables 1-10 attached as a single excel file with multiple tabs, with the first sheet containing column descriptions.

**Supplementary Table 1** | **Curated Dataset of LR Interactions, Pathway Associations, and Molecular Annotations for Intra- and Inter-Species Communication from the CellChat v2 Database**

This table compiles curated ligand-receptor pairs with pathway, structural, and localization annotations to support cell-cell communication analysis in spatial and single-cell transcriptomics. Each row includes: job_name (unique identifier), ligand_list/receptor_list (ligands and receptors in the interaction), pathway_name (associated signaling pathway), agonist/antagonist roles, co-receptor information, supporting evidence, functional annotation, is_neurotransmitter flag, protein family, localization, keywords, secreted/transmembrane status, surfaceome/adhesome classification, database version, ligand_type/receptor_type (species reference), additional agonist/antagonist/co-receptor lists, found_protein status, count_tokens, selected_interaction flag, and duplicate/removal indicators. Entries integrate CellChatDB (v1/v2) and manual curation, enabling systematic evaluation of signaling interactions across mouse and human contexts.

**Supplementary Table 2** | **Assessment of LR Interactions Using Docking and Interface Scoring Features (Excluding ipSAE Metrics)**

This table reports large-scale structure-based docking results and interaction quality assessments for predicted ligand-receptor LR pairs. Each entry represents a unique docking model, including job_name/model number (interaction identifiers), pathway_name (signaling context), ligand_type/receptor_type and ligand_list/receptor_list (species and protein composition), sequence statistics, interface contact counts, physicochemical interaction breakdown, binding affinity/Kd estimates, AlphaFold3 confidence metrics, and docking quality scores. Additional fields capture interface residue counts, structural composition, hydrogen bonding patterns, and energetic/buried surface area measurements.

**Supplementary Table 3** | **Assessment of LR Interactions Using ipSAE and Related Scores**

This table reports residue- and chain-level ipSAE metrics and related docking quality scores for predicted LR complexes. Each row represents a unique chain-chain interface within a predicted model, including job_name/model_id (prediction identifiers), Chn1/Chn2 (interacting chain IDs), PAE/Dist thresholds, interaction Type, calculated ipSAE values, AlphaFold-derived ipTM scores, docking quality (pDockQ, pDockQ2), local interaction confidence (LIS), normalization parameters (n0res, n0chn, n0dom), reference thresholds (d0res, d0chn, d0dom), and residue counts under interface and threshold conditions (nres1/nres2, dist1/dist2).

**Supplementary Table 4** | **Structural, biophysical, and interface scoring features for the selected one-to-one LR interactions**

This table provides predicted structural, energetic, and interface quality metrics for ligand-receptor models comprising a single chain for both ligand and receptor. Each entry includes job_name/model_id (prediction identifiers), pathway_name, ligand/receptor source (ligand_type/receptor_type), protein lists and Entrez IDs, interface alignment metrics (ipSAE variants), global quality scores (ipTM, pDockQ, LIS), residue and threshold normalization values, contact counts and physicochemical breakdown, binding affinity/Kd, AlphaFold3-derived disorder and confidence scores, and interface structural/energetic properties (hydrophobicity, SASA, hydrogen bonds, secondary structure). These features support detailed comparative evaluation of one-to-one LR models.

**Supplementary Table 5** | **Experimental structures and stoichiometry ratios for LR interactions**

This table compiles non-redundant experimentally resolved ligand-receptor (LR) complexes involving human and mouse proteins, including stoichiometry ratios, post-translational modifications, and sequence coverage information. Experimental data were obtained from Protein Data Bank (PDB) assemblies and curated entries from the Complex Portal. Each row includes job_name (complex identifier), pathway_name, ligand/receptor classifications, protein names and accessions, structural identifiers, stoichiometries, PTMs, source organisms, sequence ranges and coverages, and the original data source.

**Supplementary Table 6** | **Literature-supported LR interactions across and within species identified in this study**.

This table compiles ligand-receptor pairs with experimental evidence from published literature, including cross-species and same-species interactions. Each entry reports job_name (pair identifier), pathway_name, molecular classifications (ligand_class/receptor_class), species of origin, protein composition, pair_type (same vs. different species), and a Binding flag for experimentally validated interactions. Literature notes summarize structural and functional insights such as proteolytic processing, sequence conservation, post-translational modifications, and accessory proteins. Each record is linked to relevant publications (DOI) and experimentally determined or modeled structures (PDB), providing empirical context for interpreting LR predictions.

**Supplementary Table 7** | **Follow-up ligand-receptor models remodeled using alternative stoichiometry and post-translational modifications included in manuscript figures**

This table contains LR models that were re-generated to incorporate alternative stoichiometric configurations and post-translational modifications, used to illustrate structural and functional variations in manuscript figures. Each row corresponds to a remodeled model, reporting the same structural and interface quality metrics as *Supplementary Table 3*, with an additional figure_data column specifying the manuscript figure(s) in which the model appears.

**Supplementary Table 8** | **Outlier LR interactions across species**

This table lists ligand-receptor interactions exhibiting species-specific binding outlier patterns, based on comparative ipSAE analysis. For each LR pair, same-species and cross-species docking results are compared, with **delta_ipSAE** and statistical measures (**z_score, mod_z_score**) identifying extreme deviations. Additional annotations include pathway association, docking job IDs, protein lists, sequence identity, interface quality averages, and outlier classifications. Literature references, biological context, and structural flags provide functional interpretation of the observed differences.

**Supplementary Table 9** | **Amino Acid Sequences and Annotations for Selected Mouse and Human Proteins**

This table provides curated amino acid sequences and associated annotations for proteins relevant to LR dataset **(Supplementary Table 1)** in mouse (mm10) and human (GRCh38). Each row includes: the common protein name, Entrez_protein_id (NCBI identifier), ID (RefSeq or UniProt accession), full sequence, standardized name, functional description, DBxRef (database cross-references), gene name, and species of origin.

**Supplementary Table 10** | **ipSAE delta between inter- and intra-species pairs sharing a common ligand or receptor**

This table compares ipSAE values for LR pairs within species (intra-species) and between species (inter-species), restricted to cases sharing a common ligand or receptor. Each entry includes LR pair identifiers, pathway context, species types, protein composition, ipSAE values for both contexts, and the calculated delta. Unlike *Supplementary Table 8*, this table excludes annotation and literature reference columns, focusing solely on structural comparison metrics.

## References

1. Liu, Y. et al. Patient-derived xenograft models in cancer therapy: technologies and applications. Signal Transduct Target Ther 8, 160 (2023).

2. Sharma, P., Aaroe, A., Liang, J. & Puduvalli, V. K. Tumor microenvironment in glioblastoma: Current and emerging concepts. Neurooncol Adv 5, vdad009 (2023).

3. Qi, J., Luo, Z., Li, C. Y., Wang, J. & Ding, W. Interpretable niche-based cell–cell communication inference using multi-view graph neural networks. Nature Computational Science 2025 1-12 (2025) doi:10.1038/s43588-025-00809-6.

4. Jin, S., Plikus, M. V. & Nie, Q. CellChat for systematic analysis of cell-cell communication from single-cell transcriptomics. Nature Protocols 2024 20:1 20, 180–219 (2024).

5. Efremova, M., Vento-Tormo, M., Teichmann, S. A. & Vento-Tormo, R. CellPhoneDB: inferring cell-cell communication from combined expression of multi-subunit ligand-receptor complexes. Nat Protoc (2020) doi:10.1038/s41596-020-0292-x.

6. Raredon, M. S. B. et al. Comprehensive visualization of cell-cell interactions in single-cell and spatial transcriptomics with NICHES. Bioinformatics 39, (2023).

7. Manoharan, V. T. et al. Spatiotemporal modeling reveals high-resolution invasion states in glioblastoma. Genome Biology 2024 25:1 25, 1–32 (2024).

8. Liang, W. C. et al. Cross-species vascular endothelial growth factor (VEGF)-blocking antibodies completely inhibit the growth of human tumor xenografts and measure the contribution of stromal VEGF. Journal of Biological Chemistry 281, 951–961 (2006).

9. Asfaha, S. et al. Mice That Express Human Interleukin-8 Have Increased Mobilization of Immature Myeloid Cells, Which Exacerbates Inflammation and Accelerates Colon Carcinogenesis. Gastroenterology 144, 155 (2012).

10. Schwartz, A. S., Yu, J., Gardenour, K. R., Finley, R. L. & Ideker, T. Cost effective strategies for completing the Interactome. Nat Methods 6, 55 (2008).

11. Abramson, J. et al. Accurate structure prediction of biomolecular interactions with AlphaFold 3. Nature 2024 630:8016 630, 493–500 (2024).

12. Homma, F., Huang, J. & van der Hoorn, R. A. L. AlphaFold-Multimer predicts cross-kingdom interactions at the plant-pathogen interface. Nat Commun 14, 6040 (2023).

13. Dunbrack, R. L. Rēs ipSAE loquunt: What’s wrong with AlphaFold’s ipTM score and how to fix it. bioRxiv (2025) doi:10.1101/2025.02.10.637595.

14. Xue, L. C., Rodrigues, J. P., Kastritis, P. L., Bonvin, A. M. & Vangone, A. PRODIGY: a web server for predicting the binding affinity of protein-protein complexes. Bioinformatics 32, 3676–3678 (2016).

15. Pacesa, M. et al. BindCraft: one-shot design of functional protein binders. BioRxiv 2024.09.30.615802 (2024) doi:10.1101/2024.09.30.615802.

16. Bryant, P., Pozzati, G. & Elofsson, A. Improved prediction of protein-protein interactions using AlphaFold2. Nature Communications 2022 13:1 13, 1–11 (2022).

17. Zhu, W., Shenoy, A., Kundrotas, P. & Elofsson, A. Evaluation of AlphaFold-Multimer prediction on multi-chain protein complexes. Bioinformatics 39, (2023).

18. Paddock, C., Zhou, D., Lertkiatmongkol, P., Newman, P. J. & Zhu, J. Structural basis for PECAM-1 homophilic binding. Blood 127, 1052 (2015).

19. Gilbreth, R. N. et al. Crystal structure of the human 4-1BB/4-1BBL complex. Journal of Biological Chemistry 293, 9880–9891 (2018).

20. Bitra, A., Doukov, T., Destito, G., Croft, M. & Zajonc, D. M. Crystal structure of the m4-1BB/4-1BBL complex reveals an unusual dimeric ligand that undergoes structural changes upon 4-1BB receptor binding. J Biol Chem 294, 1831 (2018).

21. Wang, F. et al. Structures of mouse and human GITR-GITRL complexes reveal unique TNF superfamily interactions. Nature Communications 2021 12:1 12, 1–9 (2021).

22. Francis, B. H., Baskin, D. G., Saunders, D. R. & Ensinck, J. W. Distribution of somatostatin-14 and somatostatin-28 gastrointestinal-pancreatic cells of rats and humans. Gastroenterology 99, 1283–1291 (1990).

23. Islam, S. A. et al. Mouse CCL8, a CCR8 agonist, promotes atopic dermatitis by recruiting IL-5+ TH2 cells. Nature Immunology 2011 12:2 12, 167–177 (2011).

24. Bossen, C. et al. Interactions of tumor necrosis factor (TNF) and TNF receptor family members in the mouse and human. Journal of Biological Chemistry 281, 13964–13971 (2006).

25. Liu, J., Neupane, P. & Cheng, J. Accurate Prediction of Protein Complex Stoichiometry by Integrating AlphaFold3 and Template Information. bioRxiv 2025.01.12.632663 (2025) doi:10.1101/2025.01.12.632663.

26. Liu, X. et al. Specific Regulation of NRG1 Isoform Expression by Neuronal Activity. The Journal of Neuroscience 31, 8491 (2011).

27. Zhang, Z. et al. Tumor Microenvironment-Derived NRG1 Promotes Antiandrogen Resistance in Prostate Cancer. Cancer Cell 38, 279-296.e9 (2020).

28. Browaeys, R., Saelens, W. & Saeys, Y. NicheNet: modeling intercellular communication by linking ligands to target genes. Nature Methods 2019 17:2 17, 159–162 (2019).

29. Liberzon, A. et al. The Molecular Signatures Database Hallmark Gene Set Collection. Cell Syst 1, 417–425 (2015).

30. Kim, A.-R. et al. Enhanced Protein-Protein Interaction Discovery via AlphaFold-Multimer. bioRxiv 2024.02.19.580970 (2024) doi:10.1101/2024.02.19.580970.

31. Jumper, J. et al. Highly accurate protein structure prediction with AlphaFold. Nature 2021 596:7873 596, 583–589 (2021).

32. Virtanen, P. et al. SciPy 1.0: fundamental algorithms for scientific computing in Python. Nat Methods 17, 261–272 (2020).

33. Cock, P. J. A. et al. Biopython: freely available Python tools for computational molecular biology and bioinformatics. Bioinformatics 25, 1422–1423 (2009).

34. Bateman, A. et al. UniProt: the Universal Protein Knowledgebase in 2025. Nucleic Acids Res 53, D609–D617 (2025).

35. Beisser, D., Klau, G. W., Dandekar, T., Müller, T. & Dittrich, M. T. BioNet: an R-Package for the functional analysis of biological networks. Bioinformatics 26, 1129–1130 (2010).

36. Burley, S. K. et al. Updated resources for exploring experimentally-determined PDB structures and Computed Structure Models at the RCSB Protein Data Bank. Nucleic Acids Res 53, D564–D574 (2025).

